# Prediction of future input explains lateral connectivity in primary visual cortex

**DOI:** 10.1101/2024.05.29.594076

**Authors:** Sebastian Klavinskis-Whiting, Emil Fristed, Yosef Singer, M Florencia Iacaruso, Andrew J King, Nicol S Harper

## Abstract

Neurons in primary visual cortex (V1) show a remarkable functional specificity in their pre- and postsynaptic partners. Recent work has revealed a variety of wiring biases describing how the short- and long-range connections of V1 neurons relate to their tuning properties. However, it is less clear whether these connectivity rules are based on some underlying principle of cortical organization. Here, we show that the functional specificity of V1 connections emerges naturally in a recurrent neural network optimized to predict upcoming sensory inputs for natural visual stimuli. This temporal prediction model reproduces the complex relationships between the connectivity of V1 neurons and their orientation and direction preferences, the tendency of highly connected neurons to respond more similarly to natural movies, and differences in the functional connectivity of excitatory and inhibitory V1 populations. Together, these findings provide a principled explanation for the functional and anatomical properties of early sensory cortex.

## Introduction

An increasing number of studies – mostly focusing on mouse primary visual cortex (V1) – have begun to uncover the underlying rules specifying how cortical neurons connect^1–4^. Some findings, such as the tendency of V1 neurons to synapse with other neurons that show similar orientation selectivity, follow a simple like-for-like pattern^2– 4^. In contrast, other results, such as the spatial organization of synaptic inputs to orientation- and direction-tuned visual neurons, appear more complex and less amenable to a unifying theoretical explanation^1,5^. An outstanding question, then, is how to understand these putative connectivity rules and whether they can be explained by a single general principle.

By taking a normative approach, we can ask whether the patterns of structure and function observed at the level of individual neurons are optimized for achieving a particular goal that is likely to be important behaviorally or from an evolutionary perspective. One such promising normative framework is that of temporal prediction, which posits that sensory systems are optimized to represent those features in the recent past which are predictive of the immediate future sensory inputs^6,7^. Why might temporal prediction serve as a useful objective for an organism’s sensory systems? First, if sensory systems construct a model of the world, then a good model should predict future inputs well^8,9^. Second, predictive features are valuable for both guiding actions and compensating for neural conduction and processing delays^10^, enabling, for example, a cat to catch a bird in flight. Third, extracting predictive features reduces the vast amount of information the brain needs to manage. Finally, temporal prediction requires no explicit teaching signal beyond the sensory input itself, making it inherently more biologically plausible as an unsupervised principle than supervised counterparts^11,12^.

When optimized for temporal prediction, feedforward networks have been shown to capture many of the receptive field characteristics and response properties of V1 neurons, as well as motion processing across the visual pathway^6,7^. Nevertheless, existing feedforward temporal prediction models neglect the role of recurrency, which experimental and theoretical studies have implicated in a range of key brain functions^13–15^. Here, we show that a recurrent network optimized for temporal prediction on dynamic natural visual scenes can capture many motifs of local connectivity in visual cortex. Furthermore, when we compared network models optimized for different normative objectives, temporal prediction stood out in its capacity to explain these connectivity motifs. Hence, the relationship between the connectivity patterns of V1 neurons and their response characteristics appears to be optimized to support the predictive processing of dynamic stimuli.

## Results

### Model response properties

We trained a recurrent network model to predict the upcoming visual input (40 ms ahead, consistent with the response latencies of mouse V1 neurons^16^) based on the recent stimulus history (Fig. 1A). The model was trained on a diverse dataset of bandpass-filtered movies of natural scenes, including wildlife and panning over natural environments^17^. Because the focus of this study is on the objective that cortex might be optimized for, rather than how whether that objective is hard-wired or how it is learned, we trained the network with back-propagation. After training, we first compared model unit response properties to those of neurons in mouse V1.

**Figure 1.**
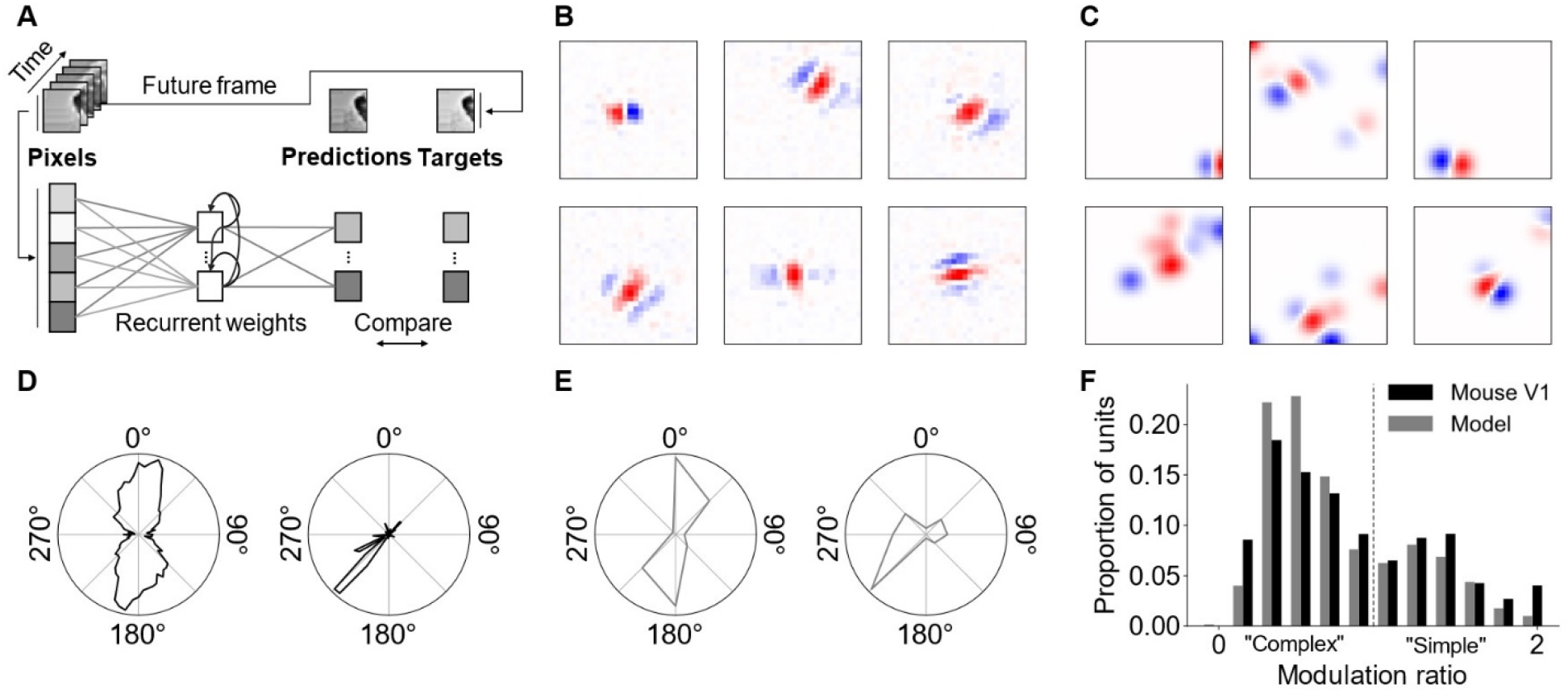
The recurrent temporal prediction model captures basic tuning properties of V1 neurons. **(A)** Schematic of the recurrent temporal model. 2,332 hidden units (90%) were excitatory with non-negative outgoing recurrent weights, the remaining 260 hidden units (10%) were inhibitory with non-positive outgoing recurrent weights. **(B)** Response-weighted-average receptive field estimates of model units. **(C)** V1 mouse receptive fields from publicly available recordings^22^, pre-processed for visualization by thresholding and smoothing with a Gaussian filter. **(D)** Exemplar tuning curves for orientation (left) and direction (right) tuned model units. **(E)** Exemplar tuning curves for orientation and direction tuned cells in V1^22^. **(F)** Distribution of modulation values across model and pooled excitatory and inhibitory mouse V1 units^21^; typically a modulation ratio <1 is taken as a complex cell and >1 as a simple cell.

We estimated the model units’ receptive fields by means of the response-weighted average for Gaussian noise movies (Fig. 1B). Like simple cells in V1 (Fig. 1C), model units often had well-defined receptive fields with a Gabor-like structure consisting of alternating, oriented excitatory and inhibitory subfields. To probe the tuning properties of the model units, we recorded the model’s response to oriented full-field drifting gratings. Model units were generally orientation tuned (24%) or direction tuned (57%; Fig. 1D, E), with a similar distribution of orientation and direction selectivity indices as found in mouse V1 (Supplementary Fig. 1). As in visual cortex, model units varied in their phase responsiveness, which was characterized by their modulation index (Fig. 1F). The responses of those units that were highly modulated by the drifting grating were classed as simple-cell-like, while units that displayed little or no phase modulation were classed as complex-cell-like^18,19^. At the population level, the model displayed a bimodal distribution similar to that found in mouse V1^20,21^, indicating the existence of distinct populations of simple-cell and complex-cell-like model units (Fig. 1F).

### Short- and long-range functional connectivity

Short-range functional connectivity between excitatory units in the model resembled that of mouse V1 neurons (Fig. 2A)^1,3^. Model units were more likely to synapse with other units with the same orientation preference (Fig. 2B), with the connection probability monotonically decreasing as the difference in orientation preference increased (*p*<0.0001, Cochran-Armitage). Likewise, connectivity between direction-tuned excitatory model units resembled that of mouse V1 neurons^3^. Model units were most likely to connect when tuned to either the same or opposite direction of motion (both *p*<0.0001, Cochran-Armitage), with connection probability decreasing as the presynaptic unit’s tuning became more orthogonal to the postsynaptic unit’s preferred direction (Fig. 2C).

**Figure 2.**
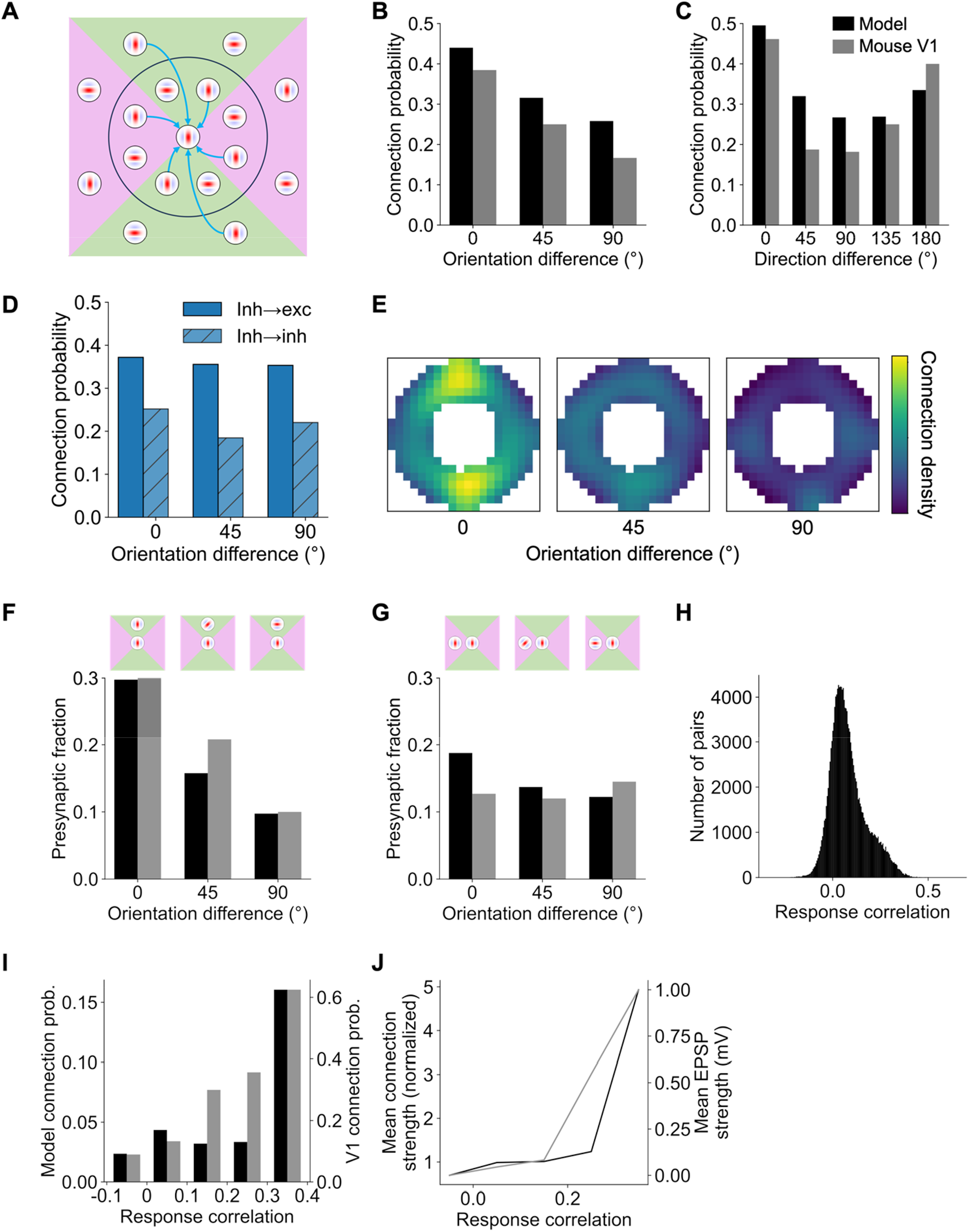
The model captures short- and long-range functional connectivity in V1. **(A)** Schematic of short- and long-range connectivity in V1. Short-range connections are more prevalent for similarly tuned V1 units, whereas long-range connection probability is greater for similarly tuned V1 neurons when their receptive fields are located in co-axial space. **(B, C)** Short-range connections are more prevalent when excitatory model units have similar orientation tuning (B) and for direction-tuned units that have similar or opposite preferred directions of motion (C), as is also the case in V1^3^. **(D)** As in B, but for inhibitory-to-inhibitory and inhibitory-to-excitatory connections in the model. **(E-G)** In both the model and V1^1^, long-range connection probability is higher for presynaptic model units with similar orientation preferences when their receptive fields are located in co-axial (F) than in co-orthogonal (G) locations relative to the receptive field of the post-synaptic unit. For B,C, F and G, data are binned using the same convention as Ko et al.^3^ with orientation bins of 0-22.5°, 22.5-67.5° and 67.5-90° and motion direction bins of 0-22.5°, 22.5-67.5°, etc. Heatmap (E) shows the normalized connection probability over visual space across differences in orientation tuning for model units. Heatmap is smoothed for display purposes with a Gaussian filter (σ=2 pixels). **(H)** Histogram of the response correlation distribution across pairs of connected model units for natural stimuli. The distribution is right skewed, indicating that a minority of model units have highly correlated responses. **(I, J)** As for mouse V1^2^, response correlation for model units co-varies with the connection probability (I) as well as the input connection strength (J). These results were abolished after randomly shuffling the recurrent weights between the units when measuring connectivity, resulting in uniform distributions (Supplementary Fig. 3). Accordingly, the model connectivity biases cannot be explained by the underlying distribution of orientation and direction tuning preferences among model units.

For inhibitory model units, co-tuning with the post-synaptic unit was much weaker (Fig. 2D), as has been previously reported in V1^5,23,24^. Neither inhibitory-to-excitatory (*p*=0.102, Cochran-Armitage) nor inhibitory-to-inhibitory (*p*=0.606, Cochran-Armitage) model unit connections showed a significant linear dependence of connection probability on the difference in orientation preference. For direction-tuned inhibitory-to-excitatory model units, the model predicts a weak but significant monotonic trend of increasing connection probability as the difference in preferred direction increases (Supplementary Fig. 2; *p*=0.032, Cochran-Armitage), distinct from the u-shaped trend observed for both excitatory model units and V1 neurons. No significant trend was found for inhibitory-to-inhibitory direction-tuned model units (*p*=0.484, Cochran-Armitage). Finally, for orientation- and direction-tuned excitatory-to-inhibitory model units, the model predicts a similar trend as for excitatory-to-excitatory model units and V1 neurons (p<0.0001, Cochran-Armitage), albeit with the minimum connection probability shifted to the 135° bin for direction-tuned model units (Supplementary Fig. 2).

Across long-range connections, the model also replicated the dependence of connectivity on neuronal orientation preference and receptive field location to that found in visual cortex^1^. We measured the connection probability between pre- and postsynaptic model units as a function of their difference in preferred orientation and the presynaptic unit’s receptive field location in visual space relative to that of the post-synaptic unit (Fig. 2E). As for mouse V1, model units were more likely to project to the post-synaptic unit if their receptive field aligned along the axis of the post-synaptic unit’s receptive field. To quantify this effect, we divided visual space relative to the postsynaptic unit into four quadrants. Those quadrants that aligned with the postsynaptic unit’s orientation tuning were referred to as ‘co-axial’ space (green regions in Fig. 2A), while those quadrants orthogonal to the unit’s preferred orientation were referred to as ‘co-orthogonal’ space (pink regions in Fig. 2A). As with the biology, orientation-tuned model units were more likely to synapse with other units when they had similar orientation selectivity and their receptive fields were located in co-axial visual space (Fig. 2F; *p*<0.0001, permutation test). Although a similar effect was found for co-orthogonal units (Fig. 2G; *p*<0.0001, permutation test), this relationship was much weaker. In particular, there was a significantly higher proportion of model units in the 0° orientation difference bin and a significantly lower proportion in the 90° bin for receptive fields in co-axial versus co-orthogonal space (both *p*<0.001, permutation test).

Similarly for V1^2,4^, model units whose responses to natural movies were highly correlated were also more likely to be connected. In both cases, the distribution of correlation values between pairs of connected model units was skewed, with the majority of pairs of units showing a relatively low response correlation, and a smaller proportion that were much more highly correlated (Fig. 2H). A qualitatively similar relationship was seen for both model units and V1 neurons between the response correlation and both the connection probability (Fig. 2I; *p*<0.001, Cochran-Armitage) and the average strength of those connections (Fig. 2J).

### Excitatory-inhibitory functional connectivity of direction-tuned units

As for orientation-tuned cells, synaptic inputs to direction-tuned cells in V1^5^ are not uniformly distributed in visual space (Fig. 3A). In particular, direction-tuned excitatory cells preferentially receive inputs from other excitatory cells whose receptive fields are situated in the opposite half of visual space relative to the postsynaptic cell’s preferred direction. In contrast, the opposite effect is observed for inhibitory cells, which preferentially synapse with excitatory cells if the location of their receptive fields is aligned with the post-synaptic cell’s preferred direction of motion. This connectivity motif provides a plausible circuit basis for direction selectivity, where a spatial offset combined with a delay in the conductance for inhibitory cells relative to excitatory cells facilitates the detection of moving stimuli^5^.

**Figure 3.**
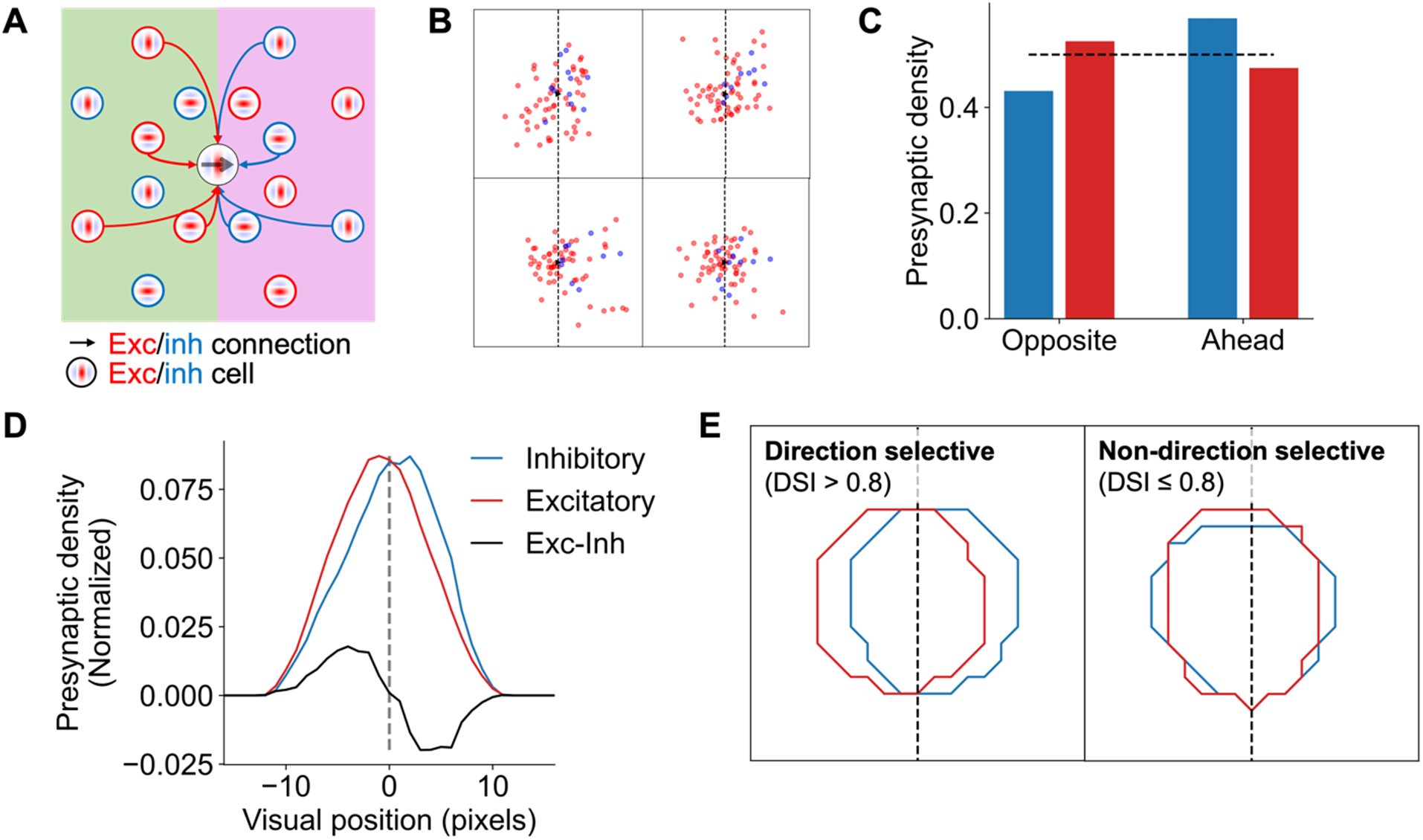
The model captures direction-dependent differences in functional connectivity between excitatory and inhibitory populations in V1. **(A)** Schematic of connectivity biases in excitatory and inhibitory inputs to direction-tuned cells in V1^5^. **(B)** Exemplar excitatory and inhibitory presynaptic ensembles for direction-tuned excitatory model units. **(C)** Model unit presynaptic density for excitatory and inhibitory cells in the halves of visual space ahead of and opposite to the postsynaptic unit’s preferred direction of motion. Dashed line represents equal density (0.5). **(D)** Profile of model unit presynaptic density across horizontal visual space for excitatory and inhibitory inputs. Profiles smoothed with a 5-pixel moving average. **(E)** Pooled density contours across all excitatory (red) and inhibitory (blue) model units for direction and non-direction selective post-synaptic excitatory units. These results were abolished after randomly shuffling the recurrent weights between the units when measuring connectivity, with no difference in density across either sector of visual space for excitatory or inhibitory model units (Supplementary Fig. 4).

In line with this evidence from V1, excitatory presynaptic ensembles in the model were more numerous in the opposite sector relative to the postsynaptic unit’s preferred motion direction (Fig. 3B-E; *t*(113)=2.93, *p*=0.004). Conversely, the opposite pattern was found for the inhibitory model units, which showed higher connection probability with direction-tuned excitatory units if their receptive fields were located in the sector of visual space ahead of the preferred direction of motion (Fig. 3B-E; *t*(113)=-2.80, *p*=0.006). This effect was specific to direction-tuned excitatory model units, with no significant difference in the spatial locations of excitatory and inhibitory presynaptic ensembles that synapse with units showing weak or no direction selectivity, as defined by a DSI ≤ 0.8 (see Methods) (Fig. 3E; *t*(113)=225, *p*=0.337).

### Comparing V1 response prediction and connectivity across models

To assess how well temporal prediction performed as a normative model of mouse visual cortex, we examined how well it captured the properties of V1 neurons relative to a series of other commonly used models across two dimensions. First, to directly compare each model’s learned representations to those in V1, we regressed the model’s hidden unit activity to predict the single-unit responses of V1 neurons to natural movies (Fig. 4A). Second, we quantitatively compared the similarity of model unit properties and their connectivity biases to those found in V1. In each of these cases, we found that the temporal prediction model both best predicted neural responses and best accounted for V1 connectivity motifs.

**Figure 4.**
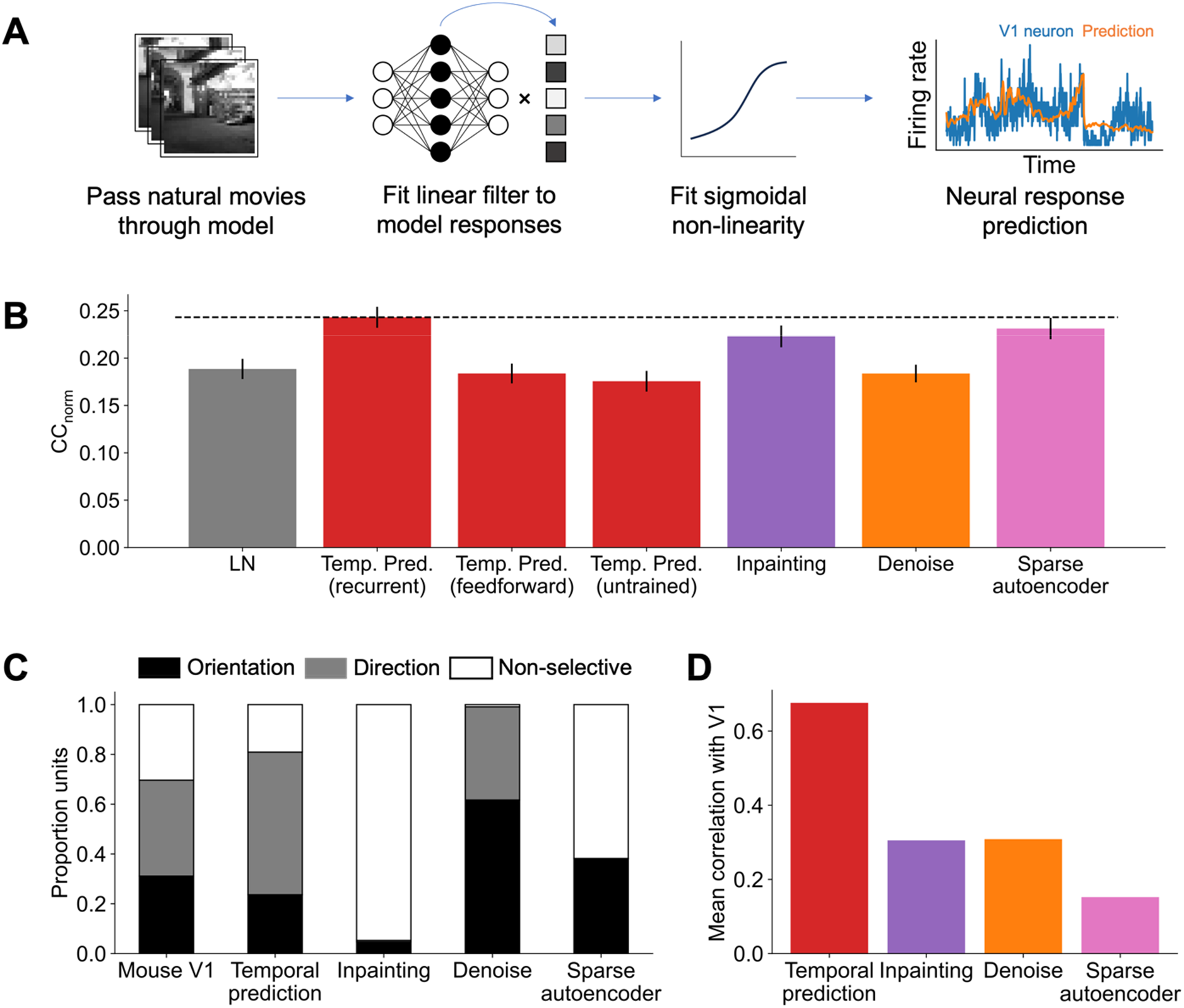
Comparison of neural prediction performance and model connectivity to V1 across unsupervised normative models. **(A)** Schematic of the neural response fitting procedure. **(B)** Performance (average CC_norm_) of the recurrent temporal prediction model (dashed line) relative to other comparison models. **(C)** Distribution of orientation-, direction- or non-selective units across mouse V1 and each model. **(D)** Comparison for each model of the average correlation of the functional connectivity profiles (Figs. 2B, C, F, G, I, J) with those found in mouse V1. A higher correlation indicates an overall better fit with the V1 data.

We compared the recurrent temporal prediction model to several other models: a linear non-linear (LN) baseline model, two variants of our temporal prediction model, as well as three models based on the same recurrent network architecture but trained using alternative learning objectives. The LN model consists of the same linear fit and sigmoidal non-linearity as for other models but is applied directly to the input stimuli. The untrained recurrent and feedforward temporal prediction networks aimed to isolate the effects of training (as opposed to model architecture alone) and recurrency. Finally, the denoising^25^, in-painting^26^ and sparse autoencoder^27^ networks all used the same architecture as the recurrent temporal prediction, but varied in their training objective. The denoising network aimed to recover the original frame from noise, the in-painting network aimed to predict the complete frame given inputs that had patches blanked out, and the sparse autoencoder was trained to reproduce the current frame while minimizing the number of internal units activated.

Neural recordings (15 recordings, *n* = 744 V1 neurons) were taken from the Allen Institute, consisting of awake mouse V1 single-unit responses to two short movie clips^22^. To predict V1 responses, each model’s hidden activity was recorded in response to the same natural movies used for the neural recordings. These responses at time *t-1* and *t-2* were then regressed to predict the neural firing rate in response to the frame at time *t*. Finally, a rectified sigmoid non-linearity was fitted to minimize the difference between the non-linear estimate of neural activity and true neural activity^7,28^. Performance was measured as the normalized correlation coefficient (CC_norm_), which describes the correlation between the predicted and true response, taking into account non-stimulus related variance in neural activity^29^.

Overall, the recurrent temporal prediction model compared favorably with the other models surveyed (mean CC_norm_=0.243, Fig. 4B). The LN model performed significantly worse (CC_norm_=0.189, *t*(743)=-6.13, *p*<0.0001, paired t-test), while the temporal prediction networks also benefited significantly from both training (untrained network mean CC_norm_=0.176, *t*(743)=-7.12, *p*<0.0001, paired t-test) and the addition of recurrency (feedforward network mean CC_norm_=0.184, *t*(743)=-5.84, *p*<0.0001, paired t-test), implying the importance of these features for modeling V1 neural responses. Finally, the recurrent temporal prediction model performed significantly better than the in-painting network (mean CC_norm_=0.223, *t*(743)=-2.10, *p*=0.036, paired t-test), the denoising network (mean CC_norm_=0.184, *t*(743)=-5.96, *p*<0.0001, paired t-test), and non-significantly better than the sparse autoencoder network (mean CC_norm_=0.231, *t*(743)=-1.27, *p*=0.204, paired t-test), again highlighting the importance of the training objective beyond model architecture alone.

The distribution of orientation- and direction-selective model units varied substantially among these models (Fig 4C). The temporal prediction model most closely reproduced the overall distribution of unit types found in mouse V1, with comparable proportions of orientation-selective (V1=31%, temporal prediction=24%), direction-selective (V1=39%, temporal prediction=57%) and non-selective units (V1=30%, temporal prediction=19%), albeit with an overrepresentation of direction-selective units. While the denoising network displayed an approximately equivalent proportion of direction-selective units (37%) to mouse V1, orientation-selective units were greatly overrepresented (62%). In contrast, no units in the inpainting or sparse autoencoder networks met the criteria for direction-selectivity, implying that these models’ learned representations did not have a temporal component.

In tandem, we compared how well each model recapitulated the connectivity biases found in mouse V1 (Fig. 4D). Specifically, for each model, we calculated a model connectivity score as the average correlation coefficient between the functional connectivity profiles of model units and V1 neurons for different response properties (Figs. 2B, C, F, G, I, J). Overall, the temporal prediction model (mean=0.68) had a much closer correspondence to the connectivity profiles found in mouse V1 compared with the other models (inpainting mean=0.31, denoising mean=0.31, sparse autoencoder=0.15). Notably, we found no significant correlation between each model’s connectivity score and its neural prediction performance (CC_norm_; *r*=0.39, *p*=0.609). Thus, the capacity of a model to predict neural responses in V1 does not imply that it can accurately capture the underlying organization of cortical connectivity.

## Discussion

The recurrent temporal prediction model exhibits response properties and functional connectivity patterns remarkably akin to those found in mouse V1, providing a unifying normative explanation for these wiring biases. In particular, the model captured the relationship between both short- and long-range connectivity patterns and neuronal tuning preferences, as well as spatial differences in the inputs from excitatory and inhibitory cells to direction-selective cortical neurons.

The extent to which cortical circuits are fundamentally stereotyped remains an enduring question in systems neuroscience. The concept of a canonical microcircuit proposes that cortical networks follow the same basic organization in which functional differences are defined primarily by their inputs and outputs, rather than by idiosyncratic, local circuits^30,31^. In support of this hypothesis, recurring cortico-thalamic and cortico-cortical loop motifs as well as cell-type- and layer-specific patterns of connectivity have been found to be consistent across many cortical areas^32,33^. However, the extent to which cortex-wide connectivity motifs might extend to the functional level is still emerging^34^.

Within mouse V1, where most work on functional connectomics has been conducted, there is growing evidence that corticocortical connections predominantly target neurons with similar feature preferences. Thus, V1 neurons are most likely to synapse with other V1 neurons that share similar orientation^1,3,35^ or direction tuning^1,3^ and exhibit correlated responses to natural stimuli^2,4^. Indeed, there is evidence that this like-for-like bias extends beyond primary visual cortex, with higher noise-correlations observed across higher-visual areas between neurons with similar tuning properties, suggesting that information is propagated both within and across cortical regions in distinct, segregated channels^36^.

In tandem with this like-for-like wiring bias, a spatial bias in local connectivity has been shown to exist for both orientation and direction tuning, whereby synaptic inputs are not uniformly distributed but rather concentrate along a specific axis. For orientation-selective neurons, connection probability is highest for similarly tuned neurons whose receptive fields are located in regions of visual space co-axial to the postsynaptic neuron’s preferred orientation^1^. Similarly, those presynaptic ensembles that synapse with direction-tuned neurons are biased to the regions of visual space ahead of (for inhibitory inputs) or opposite to (for excitatory inputs) the postsynaptic neuron’s preferred direction of motion^5,37^.

Our study demonstrates that a plausible computational principle – temporal prediction – can account for these functional connectivity patterns. Crucially, the network model was not optimized for specific response properties of visual neurons (e.g., particular receptive field characteristics). Instead, the resulting patterns of connectivity arose naturally as an emergent function of optimizing for the more general objective of predicting the neurons’ future inputs. These results suggest that wiring biases found in mouse V1 are not arbitrary but rather that they underpin an important cortical function.

### Comparison to other normative models

Despite the clear functional importance of recurrent connectivity in V1, there are comparatively few normative modeling studies addressing this topic. The key contribution of the present work is in uniting different aspects of both short-range and long-range V1 connectivity with neuronal feature preferences under a single unsupervised learning objective.

Sparse coding networks have been widely employed in modeling receptive fields and response properties^17,38,39^, and more recently have been applied to local connectivity in visual cortex. Sparse coding argues that the brain is optimized to represent stimuli efficiently such that only a small number of neurons are strongly activated at a given time. When trained on static images, sparse coding models have been shown to replicate the like-for-like connectivity pattern among units with similar orientation tuning^40^. Where motion has been included in these models, they have also been shown to capture the asymmetry in excitatory and inhibitory inputs for direction tuning^41^. However, these sparse coding models have been shown to replicate simple-cell responses only, but not complex-cell responses. Similarly, such models have not been shown to reproduce distinct connectivity profiles across separate excitatory and inhibitory cell populations, nor the long-range tuning biases reported in the present study.

Finally, local recurrent connectivity in V1 has also been approached from a Bayesian perspective, where the dependence of cortical connectivity on the similarity in orientation tuning is argued to represent an optimal means of integrating contextual information^42^. However, this Bayesian model depends on hard-coded basis functions derived from V1 simple cells and unlike our approach, cannot be said to be truly unsupervised nor learned exclusively from natural stimulus statistics.

In the context of the current study, we found that only the temporal prediction model could closely reproduce the observed relationships between V1 neurons and their functional connectivity. Thus, the results cannot be accounted for by the choice of dataset or model architecture, but are specific to the temporal prediction model’s training objective. The temporal prediction model therefore provides a more complete explanation than other models for the relationship between the connectivity of visual cortical neurons and the stimulus features to which they are tuned. In turn, these results suggest that the functional specificity of connections in V1 enables the brain to process dynamic stimuli by facilitating the prediction of upcoming sensory information. These predictions are critical for guiding complex actions^9,10^ – such as those required to catch moving prey or, in the case of a tennis player, to return the ball – which depend on estimating the future state of the world.

In systems neuroscience more widely, neural networks have recently proved to be influential in modeling sensory cortex and represent the state of the art for predicting neural responses across sensory cortex^43,44^. However, where many different models perform equivalently or near-equivalently, mere predictive power of the neural response is insufficient to identify ‘brain-like’ models. One solution to disambiguate competing models is to provide a larger space of model-brain comparisons. Notably, current approaches almost entirely neglect any correspondence in anatomical connectivity between brain and model. Indeed, as our findings illustrate, such a comparison provides scope to further distinguish between models that just have high representational capacity to predict neural responses from those that also capture underlying mechanistic similarities to the brain.

### Comparison to the biology

From an evolutionary perspective, temporal prediction is likely to confer several advantages. By encoding only those features that are predictive of future sensory inputs, temporal prediction more efficiently represents sensory data by providing a principled way of discarding non-predictive, and therefore less behaviorally relevant, information^9^. Moreover, because spiking activity constitutes the primary metabolic cost for neurons^45^, minimizing such activity through a more efficient coding scheme provides a clear adaptive advantage. Finally, given the inherent delays due to neural conduction and processing, some form of predictive processing may be essential to accurately guide motor outputs^10^.

While the current temporal prediction model is trained using backpropagation through time, the principle of temporal prediction itself is largely agnostic to the underlying learning mechanisms. Indeed, novel and more biologically-plausible learning algorithms are being developed that could, in principle, be applied to learn temporal prediction for the current network^46,47^. In this sense, the present work does not preclude either a hard-wired or learned origin for the connectivity patterns found in visual cortex^6,7^.

Given that the model’s structure is learned from an initial random state, such a configuration can, at least in theory, emerge from the interplay of some optimization principle and the natural statistics of visual inputs. Following the onset of vision, the connectivity of mouse V1 neurons that respond to similar visual features progressively increases^48^. These response-specific connectivity patterns still develop in dark-reared mice, indicating that the emergence of like-for-like wiring biases is not dependent on visual experience^49^. Nevertheless, the relationship between connection probability and the similarity of V1 responses to natural movies (but not the similarity of their orientation preferences) was found to be weaker in dark-reared mice than in animals reared with normal visual inputs. Thus, it is likely that these biases in functional connectivity result from an interplay of innate developmental programs which, at least to some extent, are later fine-tuned by sensory experience^50^.

In conclusion, we show that many aspects of functional connectivity in mouse V1 can be parsimoniously described by a single framework – temporal prediction. By optimizing a recurrent network for temporal prediction, model units naturally recapitulate both structural and functional properties of mouse visual cortex. Consequently, even seemingly disparate examples of connectivity rules may be united by a simple underlying principle of cortical organization.

## Methods

### Dataset

Training data consisted of natural wildlife videos using the same dataset as described previously^6^. Videos were taken from the repository http://www.arkive.org/species and contributed by: BBC Natural History Unit, http://www.gettyimages.co.uk/footage/bbcmotiongallery; BBC Natural History Unit & Discovery Communications Inc, http://www.bbcmotiongallery.com; Granada Wild, http://www.itnsource.com; Mark Deeble & Victoria Stone Flat Dog Productions Ltd., http://www.deeblestone.com; Getty Images, http://www.gettyimages.com; National Geographic Digital Motion, http://www.ngdigitalmotion.com. In brief, videos were converted to grayscale, bandpass filtered then down sampled to 180×180 pixels. Finally, each video was cropped into non-overlapping 36×36 pixel patches of 50 frames each, leading to a total of 40,000 clips for the training dataset and 4000 clips for the validation dataset used for hyperparameter selection. In addition, to mimic the effects of noise present in the nervous system, Gaussian noise was added to each video clip during training with a signal-to-noise ratio of 6 dB^6^.

### Network model

The model was implemented as a single-layer recurrent network with a linear readout layer to project the hidden activity to the network’s output predictions. The network’s input consisted of a 50-frame video clip, where the model was trained to predict each subsequent frame given the preceding video frames in the clip. More formally, the network receives a 1296 length vector **u**[*t*] at each time step *t*, consisting of the flattened 36×36 pixel video frame. The 2592 length hidden state vector *s*[*t*] at each time step *t* is then given by:

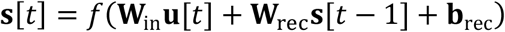

where *f* is the ReLU function, **W**_in_ is the weight matrix which describes the input weights to the network, **W**_rec_ is the weight matrix which describes the hidden, recurrent weights mapping the previous state *s*[*t* − 1] to the new hidden state *s*[*t*], and **W**_rec_ is the bias term.

The hidden activity vector *s*[*t*] at each time step *t* is then mapped to the output predictions 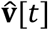 by:

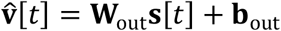

where **W**_out_ is the weight matrix describing the linear mapping from the hidden state to the output prediction and **W**_out_ is the bias term for the output weights.

In addition, to enforce Dale’s Law whereby hidden units make exclusively excitatory or inhibitory connections, each recurrent weight **W** was constrained during the forward pass as:

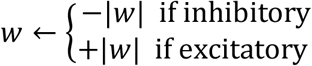

with a total of 2332 (90%) units set as excitatory and the remaining 260 (10%) as inhibitory units.

The network was then optimized using backpropagation to minimize the loss function *E*:

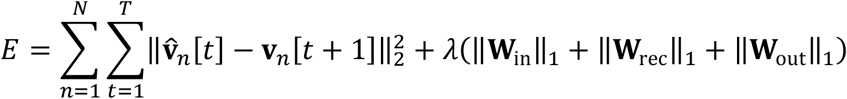

where *n* is the clip number, *N* is the total number of clips in a minibatch, *T* is the total number of time steps, and 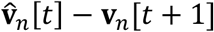 is the difference between the predicted pixel values 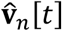 and true future pixel values ***V***_*n*_[*t* + 1]. Finally, L1 regularization is included as the sum of absolute values of all weights in the network, weighted by the λ hyperparameter.

### Implementation

The temporal prediction model was implemented in Pytorch, with gradient descent performed using the ADAM optimizer set at a learning rate of 10^−4^. The regularization strength hyperparameter *λ* was set at 10^−6^ after a hyperparameter search across lambda values (λ range = 10^-5.5^ - 10^−7^) to minimize the mean squared error on the held-out validation set.

### Comparison models

*LN model*. The LN model consisted of the same basic fitting procedure as the other models. However, where the other models regressed each network’s hidden activity, the input stimuli for the LN model were used to directly predict neural responses.

*In-painting, denoising and sparse autoencoder*. These networks consisted of the same network architecture as the recurrent temporal prediction model but with modified datasets and training objectives. For the in-painting network, the input dataset was masked with 8 randomly placed 8×8 pixel patches on each frame. For the denoising network, the input was combined with Gaussian noise with a signal-to-noise ratio of 3 dB. Finally, the sparse autoencoder was trained on the same, non-corrupted dataset as the temporal prediction model but was trained to recover the current frame under a sparsity constraint. In all these networks, the models were optimized to reproduce the current frame (rather than subsequent frame, as for the temporal prediction model) by minimizing the mean squared error between the predicted and actual current frame:

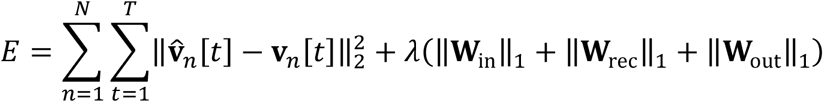

For the sparse autoencoder, an additional regularization term *λ*_act_ was included as the absolute sum of activity across all units, to encourage sparsity in the network’s representations:

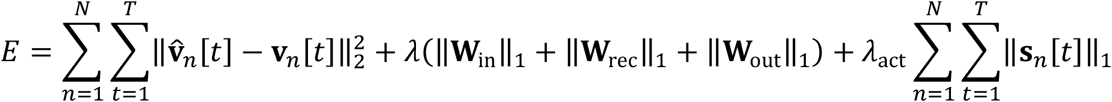

For the in-painting and denoising networks, the L1 weight regularization hyperparameter was chosen as for the temporal prediction network to minimize the mean squared error on the validation set across a range of values. For the sparse autoencoder, where no such comparable selection criterion exists, the hyperparameter set was qualitatively optimized to produce the most biologically realistic receptive fields^7^.

### Model unit analyses

*Receptive field mapping*. Model unit receptive fields were estimated using their response weighted average. In brief, the responses of model units to 25,000 frames of random Gaussian noise (μ=0, σ=1) were produced. Each noise frame was then weighted by the unit’s response to give the receptive field estimate. Model receptive fields were subsequently parameterized by a Gabor function to extract the receptive field centres and 2D extent.

#### Unit inclusion criteria

To maintain consistency across analyses, only those units whose receptive fields that were well-spatially defined and which could be well-modeled as Gabors were included for analysis. To that end, units whose receptive fields were less than 0.5 pixels in size and therefore had little spatial extent (19% of total units) or which were poorly fitted by the Gabor function were excluded (r<0.7, 12% of total units; 30% including both criteria). Short-range connections were defined as those less than 15° (2.5 pixels) and long-range as greater than 30° (5 pixels)^1^. Connections greater than 9.17 pixels were excluded to equate the experimental constraints on screen size. In the case of Ko et al.^3^, connections were not explicitly defined according to the distance between receptive fields, but we use the same short-range convention as for Iacaruso et al.^1^ that, under the assumption of retinotopy, physically short-range connections (<50 μm) are likely to be close in visual space.

### Unit tuning characteristics

To measure the model units’ tuning properties, each unit’s response to sinusoidal gratings was recorded. Gratings varied in temporal frequency (0.02-0.25 cycles/frame), spatial frequency (0.03-0.5 cycles/pixel), and orientation (0-360 degrees) with an amplitude of ±1. Each unit’s preferred temporal frequency, spatial frequency and orientation were taken as the parameter or parameter combination that maximized the unit’s mean response across 50 frames. For those analyses dependent on the unit’s spatial location, only units within the central 16×16 pixel bounds of the visual fields were included to avoid edge effects.

### Orientation and direction selectivity

Orientation and direction selectivity were quantified as OSI and DSI, respectively:

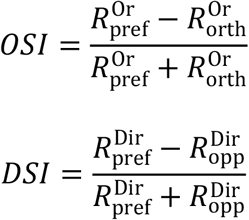

where 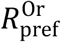 and 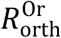 are the unit responses at the preferred and orthogonal orientations, and 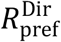 and 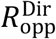 are the unit responses at the preferred and opposite (+180 degrees) directions. For Fig. 2, we take the same thresholds as Ko et al.^3^, where direction selective units defined as those with OSI values exceeding 0.4 and DSI values exceeding 0.3. For Fig. 3, where no threshold is given for Rossi et al.^5^, we take the more stringent threshold of 0.8 for direction-selective units.

#### Modulation ratio

Model units were classified as simple- or complex-like based on their phase-responsiveness to drifting grating stimuli. Quantitatively, units with a modulation ratio *F* > 1 were classed as simple-cell-like or as complex-cell-like for a modulation ratio < 1. The modulation ratio was defined as 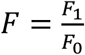 where *F*_0_ is the mean response of the neuron to its preferred stimulus and *F*_*1*_ is the amplitude of the fitted sinusoid to the neuron’s response to its preferred stimulus. Where the correlation between the fitted sinusoid and the true response was < 0.9, the modulation ratio was not further analysed for that unit.

#### Natural movie response correlations

One hundred 50-frame clips were randomly selected from the validation set and the response recorded for each model unit. The response correlation was then taken as the average correlation across the set of clips for each pair of units in the network.

### Connectivity analyses

#### Unit connectivity

Units were defined as connected if their connection strength exceeded the 95^th^ percentile of connection weights (**W**_rec_) across all pairs of units. Due to the sparse nature of the recurrent weight connectivity matrix, this threshold equated to rejecting the very low or zero weight connections, while retaining the smaller subset of highly connected units. Thus, varying this threshold across a range of values (92.5-99 percentile) did not qualitatively change the results.

#### Orientation-dependence of presynaptic organization in visual space

For each presynaptic ensemble, visual space was normalized according to the receptive field center and preferred orientation of the postsynaptic unit. Receptive field centers were first translated such that the postsynaptic unit receptive field was centered at the origin:

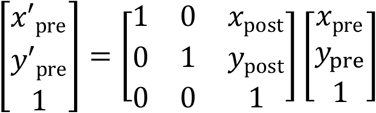

Next, receptive field centers were rotated according to the postsynaptic unit’s preferred orientation θ:

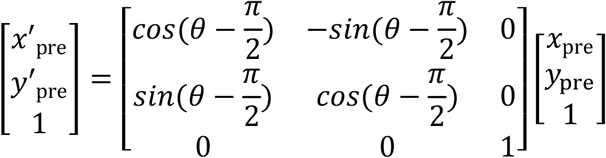

For the comparison with Iacaruso et al.^1^, presynaptic units were binned into co-orthogonal and co-axial receptive field centers according to whether they fell in one of the four quadrants orthogonal to or parallel with the postsynaptic unit’s preferred orientation (i.e., defined by *y* = −*x* and = *y* − *x*). For the comparison with Rossi et al.^5^, presynaptic units were binned according to whether they fell opposite to or ahead of the postsynaptic unit’s preferred direction of motion (i.e., defined by *x* = 0).

#### Statistical analyses

The distribution of preferred orientations and directions among model units was not uniform (Supplementary Fig. 1). To control for the possibility that the observed model connectivity distributions resulted from this overrepresentation of particular orientation and direction tuning preferences, we compared the true model results to those after randomly shuffling across model weights. Specifically, we took the total set of model units fulfilling the relevant criteria for each analysis (e.g. orientation selectivity, receptive field distance, etc.) and randomly shuffled the recurrent weights connecting these units. Note, we used these shuffled weights only for the connectivity analyses, not for any other part of any analysis, such as getting unit responses for response weighted averages. We repeated this process 1000 times, taking the mean value of the resulting distribution to compare to the true unshuffled model results.

### Neural response predictions

Neural data were taken from the Allen Institute’s Neuropixels Visual Coding dataset^22^. For each model, a linear-nonlinear mapping learned to predict the response of V1 units to natural movie stimuli (“Natural Movie One” and “Natural Movies Three”, 150 seconds total). We included all recorded V1 units from wild type mice whose noise to signal power ratio in response to the natural movies was below 60^51^.

For each model, the neural fitting process consisted of first learning a linear mapping using Lasso regression before fitting a rectified sigmoidal non-linearity^28^. Prior to fitting, the dimensionality of the model unit activity that was regressed from was reduced to the first 200 components using principal component analysis fitted on the training-set model responses. PCA was used to equate the number of parameters across models and increase the efficiency of model fitting^7^. The non-linearity 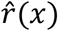 was defined as:

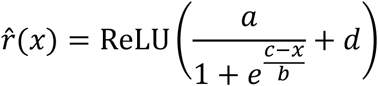

where the parameters *a, b, c* and *d* were optimized to minimize the mean squared error between the true and predicted neural firing rate using the SciPy “curve_fit” function^52^. The L1-regularization strength (α) of the Lasso chosen via cross-validation from 40 values log-spaced between 10^1^ and 10^−5^ to maximize the average CC_norm_ across each fold’s validation set for the combined linear-nonlinear mapping. The reported values are finally taken as the performance on the held-out test set.

### Mouse V1 comparisons data

Mouse data for comparison were either extracted from published figures using (Fig. 1F; Fig. 2J), taken as the exact statistics from the published paper (Fig. 2B, C, F, G, I) or computed directly from the Neuropixels Visual Coding dataset from the Allen Institute^22^ (Fig. 1C, E; Fig. 4C; Supplementary Fig. 1). For the V1 receptive fields (Fig. 1C), these were estimated by fitting a linear filter to predict the responses of V1 single units to natural movies as described above.

To compare how well each model captured the connectivity patterns described for mouse V1, we calculated a model connectivity score as the average of the Pearson correlation coefficients between the described neural connectivity profiles (data in Fig. 2B, C, F, G, I, J) and the corresponding connectivity profiles of the given model.

## Acknowledgments

This work was funded by a Wellcome Principal Research Fellowship (WT108369/Z/2015/Z) to AJK. SK-W was supported by a studentship funded by the Nuffield Department of Clinical Neurosciences at the University of Oxford.

## Code availability

All code used to generate analyses and figures presented in this manuscript will be available at https://github.com/sebbkw/temporal_prediction_connectivity.

