## Supplementary information for "Prediction of future input explains lateral connectivity in primary visual cortex"

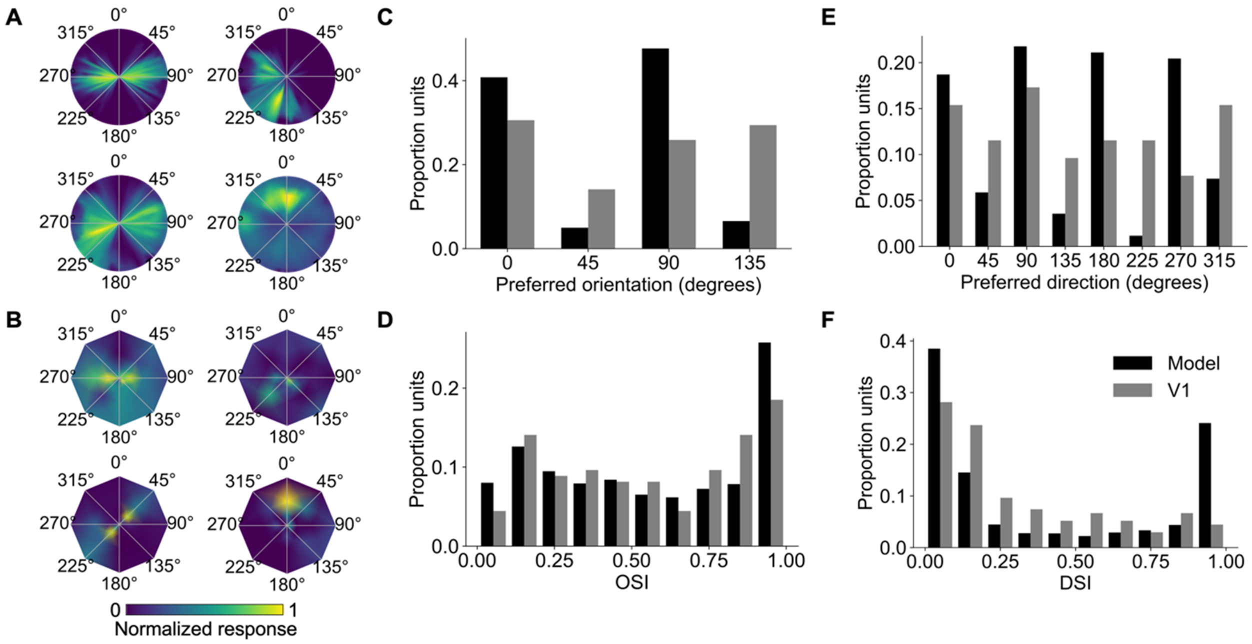


**Supplementary Figure 1. Model and mouse V1 tuning properties.** All mouse data from the Allen Brain Institute's' visual coding dataset. **(A)** Example model unit responses to drifting gratings as a function of temporal frequency (1-8 Hz, radial axis) and orientation (polar angle). **(B)** Example mouse V1 responses to drifting gratings as a function of temporal frequency (2-15 Hz, radial axis) and orientation. **(C)** Distribution of preferred drifting grating orientations for model units and mouse V1. **(D)** Distribution of orientation selectivity indices for model units and mouse V1. **(E)** Distribution of preferred drifting grating directions for model units and mouse V1. **(F)** Distribution of direction selectivity indices for model units and mouse V1.

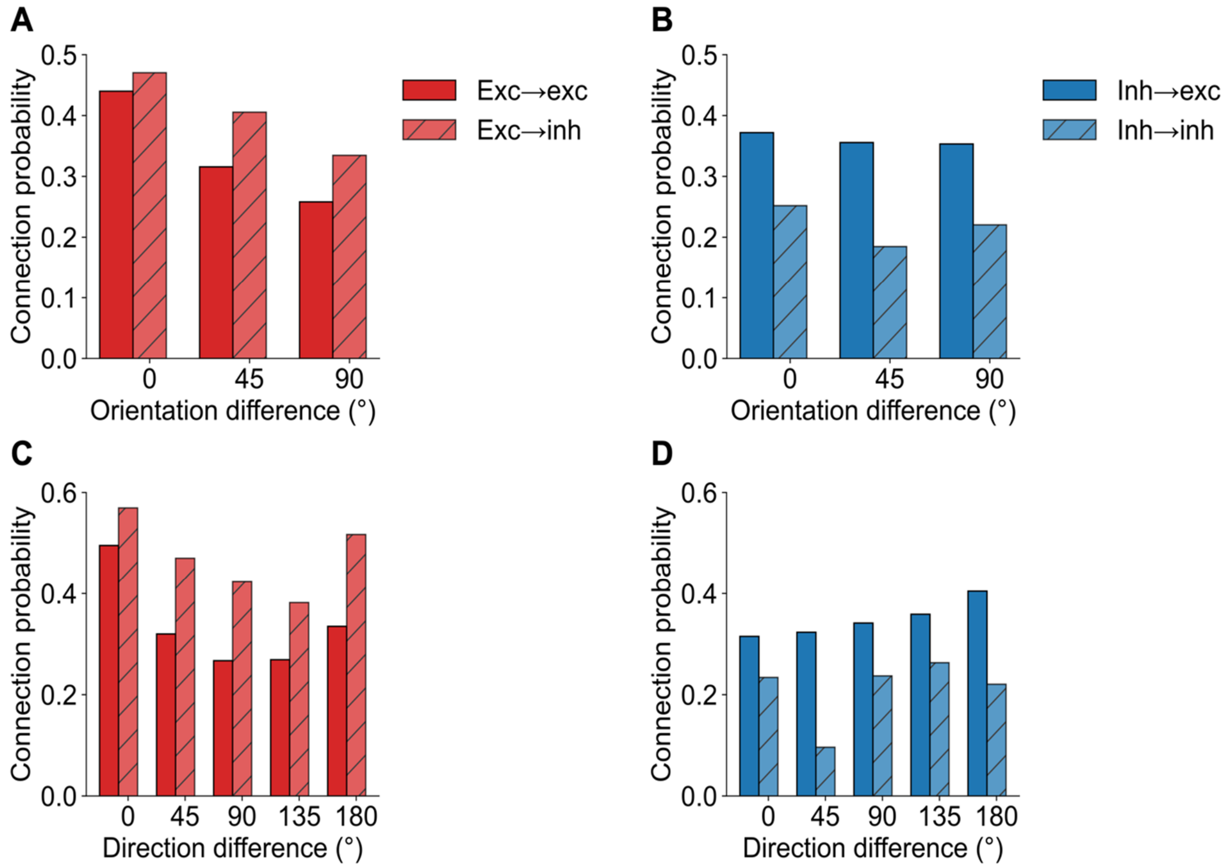


**Supplementary Figure 2. Orientation and direction dependence of connection probabilities across model unit types. (A, B)** Connection probability as a function of orientation tuning difference for excitatory (A) and inhibitory (B) model units. Excitatory units show a monotonic trend of decreasing connection probability, whereas inhibitory units show a much weaker effect, implying a broader functional spread of inhibitory connections. **(C, D)** Connection probability as a function of direction tuning difference for excitatory (C) and inhibitory (D) model units. Excitatory units show a characteristic u-shaped curve as found in V1, where similar or opposite direction-tuned units are most likely to connect. Inhibitory units synapsing with excitatory units show a weak but monotonically increasing trend, where the closer the tuning is to its opposite direction, the more likely they are to connect. Finally, inhibitory model units synapsing with other inhibitory units show a more heterogenous pattern, with no overall preference for a given difference in direction tuning.

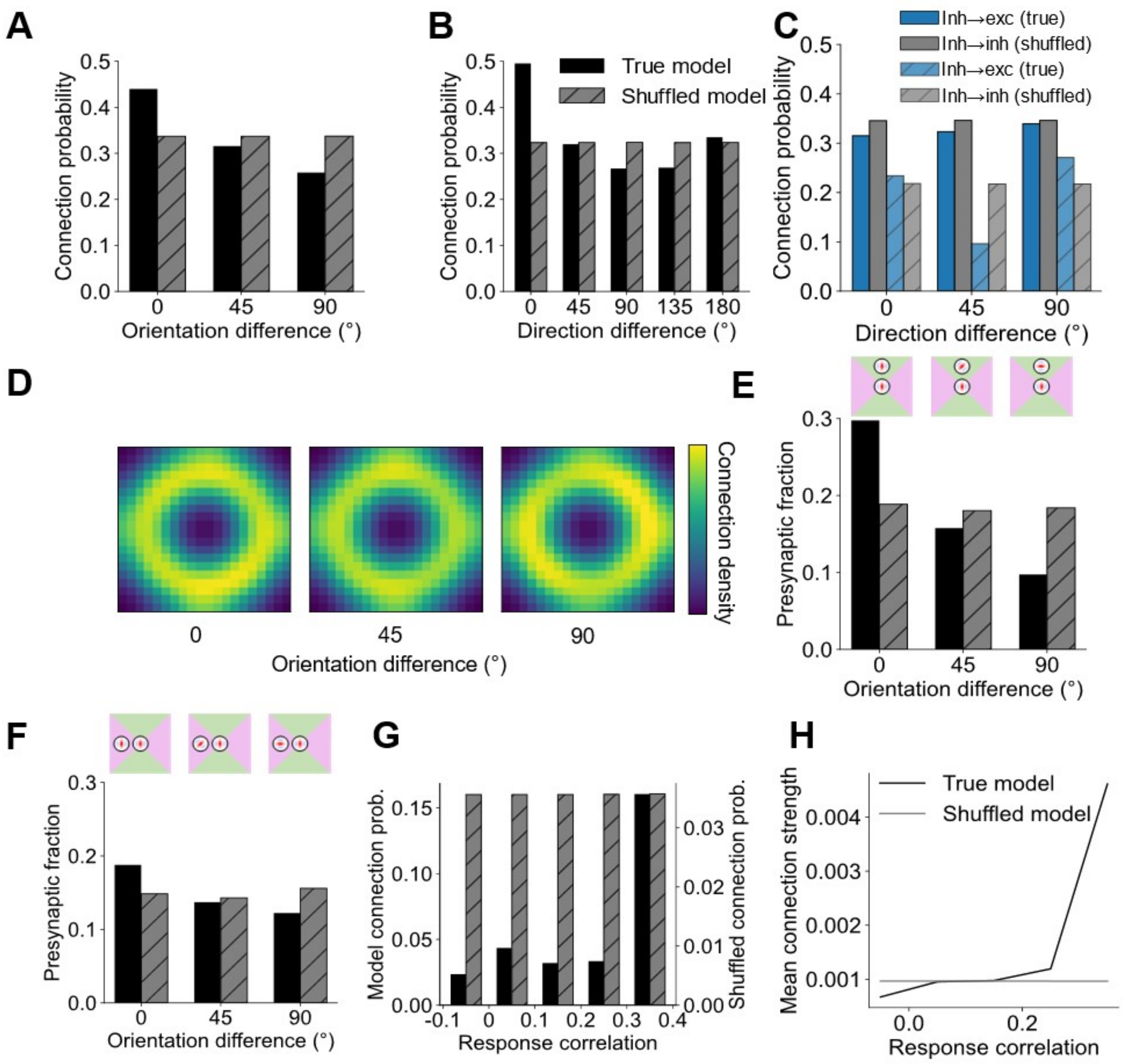


**Supplementary Figure 3. Short- and long-range functional connectivity resembling mouse V1 in the model is abolished when connectivity is measured with the recurrent weights randomly shuffled. (A, B)** Short-range connection probability as a function of the difference in orientation and direction tuning among model units. **(C)** As in A, but for inhibitory-to-inhibitory and inhibitory-to-excitatory connections in the model. **(D-F)** Long-range connection probability as a function of difference in orientation preferences for receptive fields located in co-axial (E) and co-orthogonal (F) locations relative to the receptive field of the post-synaptic unit. Heatmap (D) shows the shuffled connection probability over visual space across differences in orientation tuning for model units. Heatmap is smoothed for display purposes with a Gaussian filter (σ=2 pixels). **(G, H)** Response correlation for model units as a function of connection probability (G) as well as the input connection strength (H).

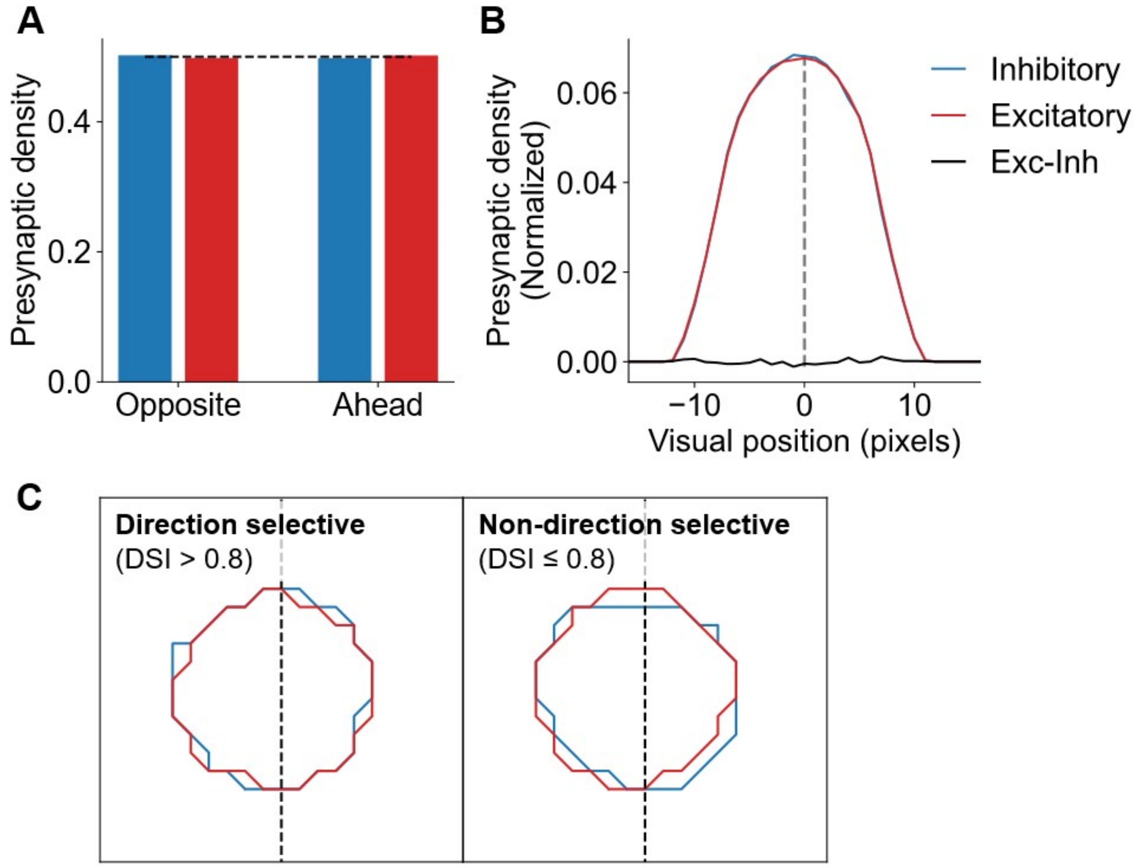


**Supplementary Figure 4. Direction-dependent differences in functional connectivity between excitatory and inhibitory populations are abolished when connectivity is measured with the recurrent weights randomly shuffled. (A)** After shuffling, model unit presynaptic density for excitatory and inhibitory cells is equal in both halves of visual space. Dashed line represents equal density (0.5). **(B)** Profile of model unit presynaptic density across horizontal visual space for excitatory and inhibitory inputs. Profiles smoothed with a 5-pixel moving average. **(C)** Pooled density contours across all excitatory (red) and inhibitory (blue) model units for direction and non-direction selective post-synaptic excitatory units, showing no overall differences after shuffling.
